# The *Dirofilaria immitis unc-49* gene encodes a pharmacologically unique cys-loop GABA receptor

**DOI:** 10.1101/2025.10.30.685166

**Authors:** Sierra Varley, Tobias Clark, Nathan Chubb, Nicolas Lamassiaude, Claude L. Charvet, Cédric Neveu, Sean G Forrester

## Abstract

Nematode cys-loop GABA receptors play important roles in movement and locomotion. GABA receptors encoded by *unc-49* appear to be widespread in nematode genomes including parasitic nematodes but are poorly characterized in filarial parasites. *Dirofilaria immitis* is a filarial parasite responsible for heartworm disease in dogs. Macrocyclic lactones are widely used to prevent heartworm infection. However, like many anthelmintics, resistance has emerged and new solutions are needed including the discovery of new anthelmintic drug targets. In this study, we report the isolation of two *unc-49* subunit mRNAs (*unc-49b* and *unc-49c*) from the canine parasitic nematode *D. immitis*. When expressed in *Xenopus* oocytes, Dim-UNC-49B formed a functional homomeric GABA-gated channel, whereas Dim-UNC-49C did not. We further demonstrated that Dim-UNC-49B and Dim-UNC-49C gave rise to a functional heteromeric channel. Pharmacological characterization of these receptors and cross-species co-expression with UNC-49 subunits from the parasitic nematode *Haemonchus contortus* showed some unique properties compared to the same receptors characterized from other nematodes. These results revealed new properties of UNC-49 receptors in filarial worms.

## Introduction

Nematode cys-loop GABA (γ-aminobutryic acid) receptors are unique types of ligand-gated chloride channels that are found at neuromuscular junctions and play important roles in movement. Compared to human GABA_A_ receptors, nematode GABA receptors share about 30-40% amino acid similarity (Bamber et al., 2003). These receptors were originally characterized *in situ* through electrophysiological investigation of the body wall muscles of the large parasitic nematode *Ascaris suum* revealing the presence of cys-loop GABA receptors at neuromuscular junctions (Martin, 1980; Martin, 1985; Holden-Dye et al., 1988; Holden-Dye et al., 1989; Holden-Dye and Walker, 1990; Martin et al., 1991). Not only are these receptors functionally unique compared to mammalian receptors but they also exhibit a unique pharmacological profile where they do not respond to the classical GABA receptor antagonist bicuculline (Holden-Dye et al. 1988, 1989). One important nematode GABA receptor is encoded by the *unc-49* gene which plays a key role in muscle contraction that is important for the sinusoidal movement of nematodes. In the free-living nematode *C. elegans, unc-49* gene mutants display a locomotion phenotype called “shrinker” due to the lack of inhibitory signals responsible for muscle relaxation (Bamber et al., 1999; McIntire et al., 1993b; McIntire et al.,1993a).

Genes encoding UNC-49 GABA receptors are found widely among both parasitic and free-living nematode species and have similar polypeptide sequences (Accardi et al 2012). However, while these receptors appear to be ubiquitous among nematodes, differences in functional properties between different nematode species have been noted. For example, there are two UNC-49 receptor subunits (B & C) that have been characterized from *C. elegans*. UNC-49B can form a functional homomeric channel that responds to GABA and can associate with UNC-49C to produce a heteromeric channel that exhibits a lower sensitivity to GABA (Bamber et al., 1999). On the other hand, in the parasitic nematode *Haemonchus contortus* the heteromeric UNC-49B/C channel has a higher sensitivity to GABA compared to the UNC-49B homomeric channel (Siddiqui et al., 2010). These differences may be the result of amino acid differences in various residues withing the GABA binding pocket (Accardi et al., 2011). This may not be surprising since there is at most an 80% homology in the UNC-49 polypeptide between *C. elegans* and *H. contortus*. However, the UNC-49 subunits in *C. elegans* and *H. contortus* do share many similar features. For instance, in both species the UNC-49B homomeric channel is highly sensitive to picrotoxin (PTX) block and the channel becomes less sensitive to PTX with the addition of UNC-49C (Siddiqui et al. 2010; Bamber et al. 2003).

Filarial nematodes, while more distantly related compared to *C. elegans* and *H. contortus*, also appear to exhibit the *unc-49* gene (Accardi et al 2012). One filarial nematode found widespread in North America and other parts of the world is the parasite *Dirofilaria immitis* which is the causative agent of heartworm disease in dogs, cats and ferrets. Heartworm infection has a serious impact on companion animals, and prevention of infection with macrocyclic lactones is the only option. One of the most pressing issues with *D. immitis* is the growing resistance to macrocyclic lactones and the lack of alternative treatment options (Geary, 2023). The mode of action of macrocyclic lactones still remains poorly understood in filarial parasite nematodes including *D. immitis*. Glutamate-gated chloride channels may mediate the preventative effects of macrocyclic lactones in *D. immitis*. Ivermectin was previously reported to potently activate the AVR-14B homomeric glutamate-gated chloride channels, which is, to date, the only functional ion channel ever characterized in *D. immitis* (Yates and Wolstenholme, 2004).

In this context, our goal was to isolate and examine the pharmacological properties of the UNC-49 GABA receptor in *D. immitis*. By utilizing a cross-species expression approach in *Xenopus laevis* oocytes we were able to observe unique functional properties of the *D. immitis* UNC-49 receptors.

## Methods

### Identification and cloning of unc-49b and unc-49c

For initial isolation of RNA, frozen adult parasites of *D. immitis* were received from Zoetis. The sample was homogenized with liquid nitrogen, and total RNA was isolated using TRIzol Reagent (Invitrogen, California, USA). Isolated RNA was stored at -80°C until used. Complementary first strand DNA (cDNA) was synthesized using the Quantitect Reverse Transcriptase kit (Qiagen, Dusseldorf, Germany), using a 3’ oligo-dT anchor primer (5’CCTCTGAAGGTTCACGGATCCACATCTAGATTTTTTTTTTTTTTTTTVN3’); where V is either A, C, or G and N is either A, C, G, or T).

Gene fragments of *unc-49* were identified from a comparative genomic screen of parasitic worms through bioinformatic analysis of the genome. From there, Basic Local Alignment Search Tool (BLAST) (Altschul et al. 1990) was used to compare sequences. NCBI was used to obtain protein sequences for UNC-49B (GenBank: ACL14329.1) and UNC-49C (GenBank: ABW22635.1) from *H. contortus*, as well as *C. elegans* UNC-49B (GenBank: AAD42384.1) and UNC-49C (GenBank: AAD42386.1). These sequences were used in tBLASTn which uses protein sequence to search a translated nucleotide database in all six reading frames (Altschul et al. 1990; Gish and States 1993). The search was limited to whole genome shotgun (wgs) contigs since it is currently the only complete genetic database available for *D. immitis* (NCBI Accession: JAKNDB000000000.1). The wgs contigs predicted by tBLASTn to contain exons coding for a protein of interest were imported into SnapGene Viewer (v6.2.1) (https://www.snapgene.com/snapgene-viewer) and translated in all 6 reading frames to identify intronic regions of the wgs contig. The predicted *unc-49* genes were pieced together based on the sequences from *H. contortus* and *C. elegans* to determine if 5’ and 3’ ends of the genes were complete with a start and stop codon and through sequence alignments. The sequence was translated *in silico* using SnapGene Viewer to ensure the complete coding sequence was being captured.

To clone the full-length genes of *unc-49b* and *c* we blasted the *D. immitis* Transcriptome Shotgun Assembly Sequence Database (TSA) to identify the full-length cDNA sequence of *dim-unc-49c* (JR913114.1). We then confirmed the 5’ ends of the sequences using 5’ RACE with SL1 [splice-leader sequence 1; (5’-GGTTTAATTACCCAAGTTTGAG-3’)] and 3’-RACE PCR. New reverse specific primers were designed to amplify and clone the full-length coding cDNAs corresponding to *dim-unc-49b* and *c*. Once it was confirmed that the complete sequence was cloned, primers were designed containing restriction sites to facilitate directional cloning into pGEMHE in the case of *dim-unc-49c* or pTB-207 in the case of *dim-unc-49b.* Once cloned the sequences were verified (Genome Quebec Montreal, Canada). Gene naming was consistent with recommendations outlined in Beech et al 2010.

### Homology Modelling and docking of ligands

The SWISS-MODEL Repository (Bienert et al. 2017; Waterhouse et al. 2018) was used to search the databases of existing three dimensional protein models against genes of interest from this study. Homology modeling of the UNC-49 receptor was done using *C. elegans* glutamate-gated chloride channel named 3RIF as a template, as determined by SWISS-MODEL. Using MODELLER 10.4 (Šali and Blundell 1993), protein sequences of interest were aligned to 3RIF in a three-dimensional conformation, which proved to be the most energetically favorable. These protein sequences were then arranged as a homodimer through MODELLER 10.4. From there, ligands were docked into the hypothetical binding pocket using AutoDockTools 1.5.7 (Trott and Olson 2010). The ligand dock with the most energetically favorable score was selected for further analysis. The visualization of the interaction of the ligand with the receptors was done using Chimera 1.16 (Pettersen et al. 2004), which also allows for putative distance calculations to be performed, and the residues in the binding pocket to be identified.

### Heterologous expression of channels in Xenopus laevis oocytes

All animal procedures followed the Ontario Tech University Animal Care Committee and the Canadian Council on Animal Care guidelines. Female frogs were supplied by NASCO (Dexter, Michigan) and housed in a temperature-controlled room with 12-hour light-dark cycling. The frogs were anesthetized with 0.15 % 3-aminobenzoic acid ethyl ester methanesulphonate salt (MS-222) (Sigma-Aldrich, Canada), and pH adjusted to 7 with sodium bicarbonate. Surgical extraction of ovaries was performed, and the oocytes were defolliculated with 2mg/ml of collagenase-II (Sigma-Aldrich, Canada) in calcium-free oocyte Ringer’s solution [82 mM NaCl, 2 mM KCl, 1 mM MgCl2, 5 mM HEPES pH 7.5 (Sigma-Aldrich, Canada)]. The oocytes were placed on a rocker and left to defolliculate at room temperature for 1 hour. The oocytes were rinsed with ND96 solution (1.8 mM CaCl_2_, 96 mM NaCl, 2 mM KCl, 1 mM MgCl_2_, 5 mM HEPES, pH 7.5) and stored in ND96 supplemented with 50 μg/mL gentamycin and 275 μg/mL of pyruvic acid (ND96-Supp) and stored at 18°C for use.

*H. contortus unc-49b* and *unc-49c* were subcloned into the pT7Ts *X. laevis* expression vector and were also used in this study (Siddiqui et al 2010). For each gene, cRNA was synthesized using the T7 mMessage mMachine kit, (Ambion, Texas, USA) and 1 μg of the linear plasmid was added to each reaction and incubated according to kit instructions. cRNA concentration and purity were assessed using a BioDrop DUO^®^, an ultraviolet–visible spectrophotometer. Samples were diluted to 400 ng/μL with Water for Molecular Biology (Sigma-Aldrich, Canada). Glass Capillaries (1.2 OD x 0.69 ID x 100 L mm) (Harvard Apparatus, Massachusetts) were pulled into needles for injection using a P-97 Micropipette Puller (Sutter Instrument Co., California, USA). The needles were backfilled with mineral oil and loaded onto a Nanoject II microinjector (Drummond Scientific, USA) and the required cRNA was loaded into the needle. For all receptor combinations tested, *X. laevis* oocytes were injected with 50 nL (20 ng) of RNA for each subunit, and oocytes were incubated at 18°C in ND96-Supp for 48 to 90 hours.

Glass electrodes (1.5OD x 0.86 ID x 100 L mm) (Harvard Apparatus, Massachusetts) were pulled using the P-97 Micropipette Puller (Sutter Instrument Co., California, USA). The electrodes were backfilled with 3 M KCL, and silver wires were inserted into the back, and the electrodes were mounted on the electrode holder. Stocks of 100 mM GABA (Sigma Aldrich, Canada) or 20 mM PTX (Sigma-Aldrich, Canada) were dissolved in ND96. Serial dilutions of GABA were made for various concentrations between 30 mM and 10 μM. Axoclamp 900A (Molecular Devices, USA) was used to perform two-electrode voltage clamp electrophysiology (TEVC) on oocytes. The oocyte was placed in the RC-1Z perfusion chamber and agonist dose response curves were generated by applying increasing concentrations of agonists over the oocyte. Data was collected by perfusing various ligands over the oocyte and measuring the current generated. Data was analyzed using Clampex Software v10.2 (Molecular Devices, USA). EC_50_ values were generated using the data from the dose-response curves using GraphPad Prism Software 8.0.2 for Windows 10 (GraphPad Software, USA).

For PTX experiments, GABA (at EC_50_ concentration) was dissolved in ND96 with increasing concentrations of PTX from 1 μM to 250 μM. For these experiments the oocytes were incubated with EC_50_ GABA concentration, followed up with a solution containing both GABA and one of the PTX experimental concentrations. This technique was repeated on a new oocyte, with an increasing amount of PTX, each time until maximum inhibition was reached. The GABA with PTX response was compared to the GABA only response for each PTX concentration tested. Inhibition response curves were generated with Graphpad Prism Software 6 (California, USA).

At least 5 oocytes, and at most 18 oocytes from 2 different frogs were used to generate each data point. Data was analyzed using students t-test in Graphpad Prism Software 6. Significance was determined if p ≤ 0.05.

## Results

### Molecular analysis of *unc-49b* and *unc-49c* from *D. immitis*

#### Characteristics of dim-unc-49b and c sequences

The cDNA transcript of *dim-unc-49b* is 1455 base pairs in length and encodes a 485 long amino acid sequence including a predicted signal peptide sequence. The protein sequence of Dim-UNC-49B had a 72% identity and an 82% similarity to Hco-UNC-49B and a 69% identity and 80% similarity to the *C. elegans* Cel-UNC-49B (supplemental A). The full-length *dim-unc-49c* was determined to be 1341 base pairs long. The protein sequence encoded by *dim-unc-49c* is 446 amino acids and contains a predicted signal peptide sequence. Dim-UNC-49C is 66% identical and 80% similar to Hco-UNC-49C and is 62% identical and 77% similar to Cel-UNC-49C (Supplemental B). As observed in other nematode species, the 5’ end of *dim-unc-49c* is identical to the 5’ end of *dim-unc-49b* (Figure 1A), and the gene only differs in its 3’ end though differential splicing (**Error! Reference source not found.**B). Binding loops D, A, E and a portion of Loop B are encoded for by the identical 5’ region in 5 exons, the remainder of the protein is encoded for by the 3’ portion which differs between the two genes (ie. Mature mRNA) variants. Exons 6 to 11 encode the UNC-49B receptor and exons 12 through 16 encode the UNC-49C receptor. Phylogenetic analysis revealed that Dim-UNC-49B and C group with the UNC-49 subunits in general but appear to more related to subunits from the *Brugia malayi* (Figure 1C).

**Figure 1.**
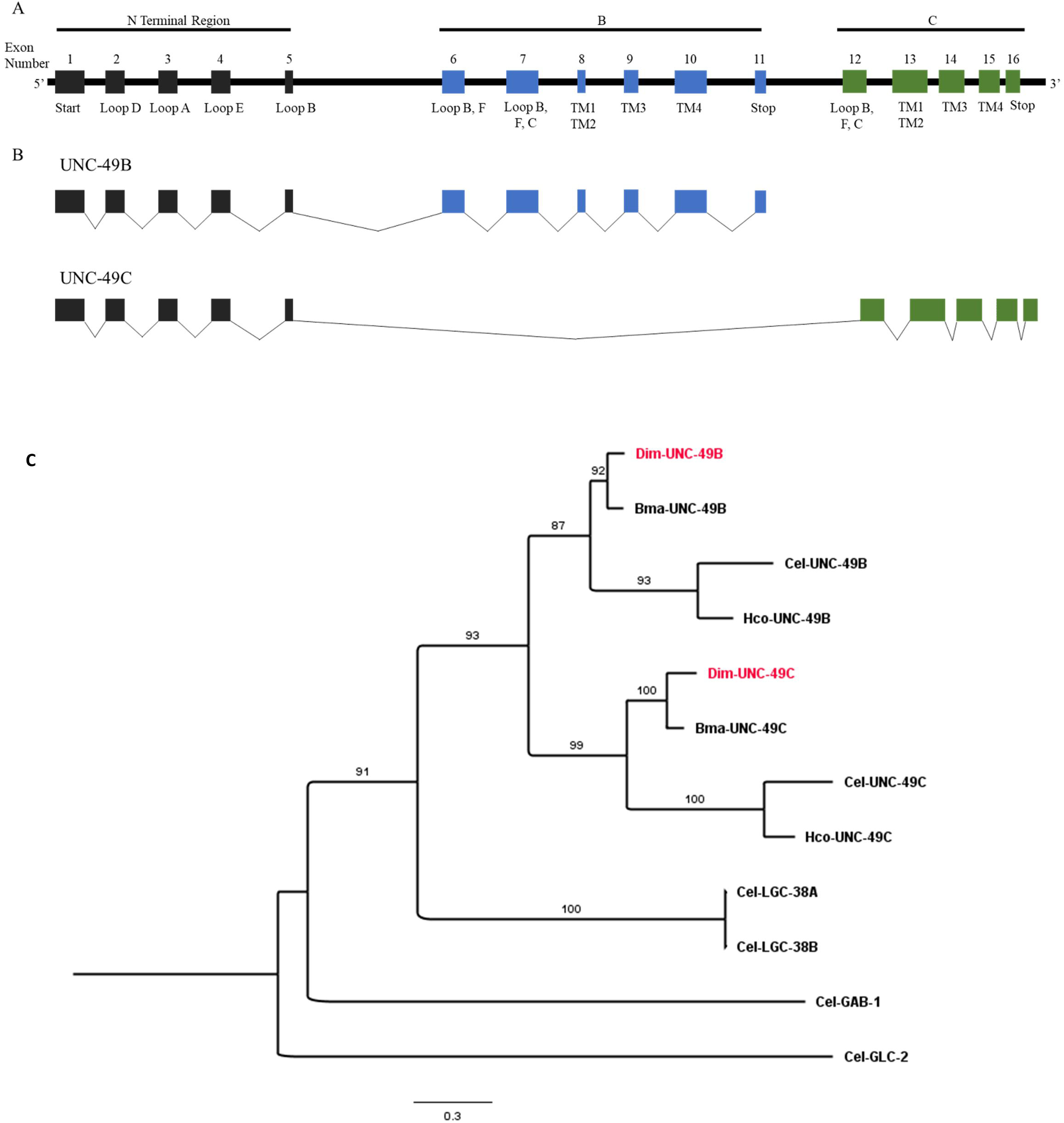
(A) Genomic organization of the *unc-49* receptors in *D. immitis* (NCBI Accession: JAKNDB000000000.1). Black indicates exons shared between both *unc-49b* and *unc-49c*. Blue indicates *unc-49b* exons, green indicates *unc-49c* exons. Binding loops are labelled, and transmembrane domains are labelled TM1-TM4 (B) Splicing pattern of the *unc-49b* and *unc-49c* genes. Black lines indicate the introns that are spliced out of the gene. (C) Maximum likelihood tree showing relationship of UNC-49 GABA receptor subunits from *Dirofilaria immitis* (Dim), *Brugia malayi* (Bma), *Haemonchus contortus* (Hco) and *Caenorhabditis elegans* (Cel) with other *C. elegans* GABA subunits. The UNC-49B and UNC-49C *D. immitis* sequences investigated in the present study are highlighted in red. The tree was built upon an alignment of complete coding sequences and rooted with the glutamate-gated chloride channel subunit GLC-2 from *C. elegans*. Boostrap values (100 replicates) are indicated on branches.

#### Sequence analysis of the major binding loops

We have shown previously that the binding pocket of the UNC-49 receptor from *H. contortus* is at the interface of two UNC-49B subunits (Accardi and Forrester 2011). The *Dirofilaria* UNC-49B subunit also contains the essential binding site residues in loops D, A, E, B and C. Interestingly, we observed an amino acid variation (M→L) in *D. immitis* loop D (Figure 2A) which has been shown to have a small impact in the GABA response in *H. contortus* UNC-49 receptors (Accardi and Forrester 2011). Sequence analysis major binding loops is shown in Figure 2A. As seen with *H. contortus* the essential binding site residues in loops D, A, E, B and C are present in the *D. immitis* UNC-49B subunit. To further visualize the interaction of GABA with binding site residues homology models were created. The model generated contains many of the predicted binding site residues that have been observed in previous models of the UNC-49 receptor from *H. contortus* and the GABA RDL receptor from *Drosophila melanogaster* (Lummis et al 2011). Specifically, the amine group of GABA is oriented towards Y190 (from loop B) and is 3.6 Å from the OH group of the tyrosine. The GABA amine group is also oriented towards Y241 (on loop C). T236 (loop C) is predicted to be 2.4 Å from the oxygen on the carboxylic acid group of GABA. On the complimentary subunit, Y87 (loop D) is a predicted 2.5 Å away from the amine group and R89 (loop D) is predicted to be 5.1 Å from GABA (Figure 2B).

**Figure 2:**
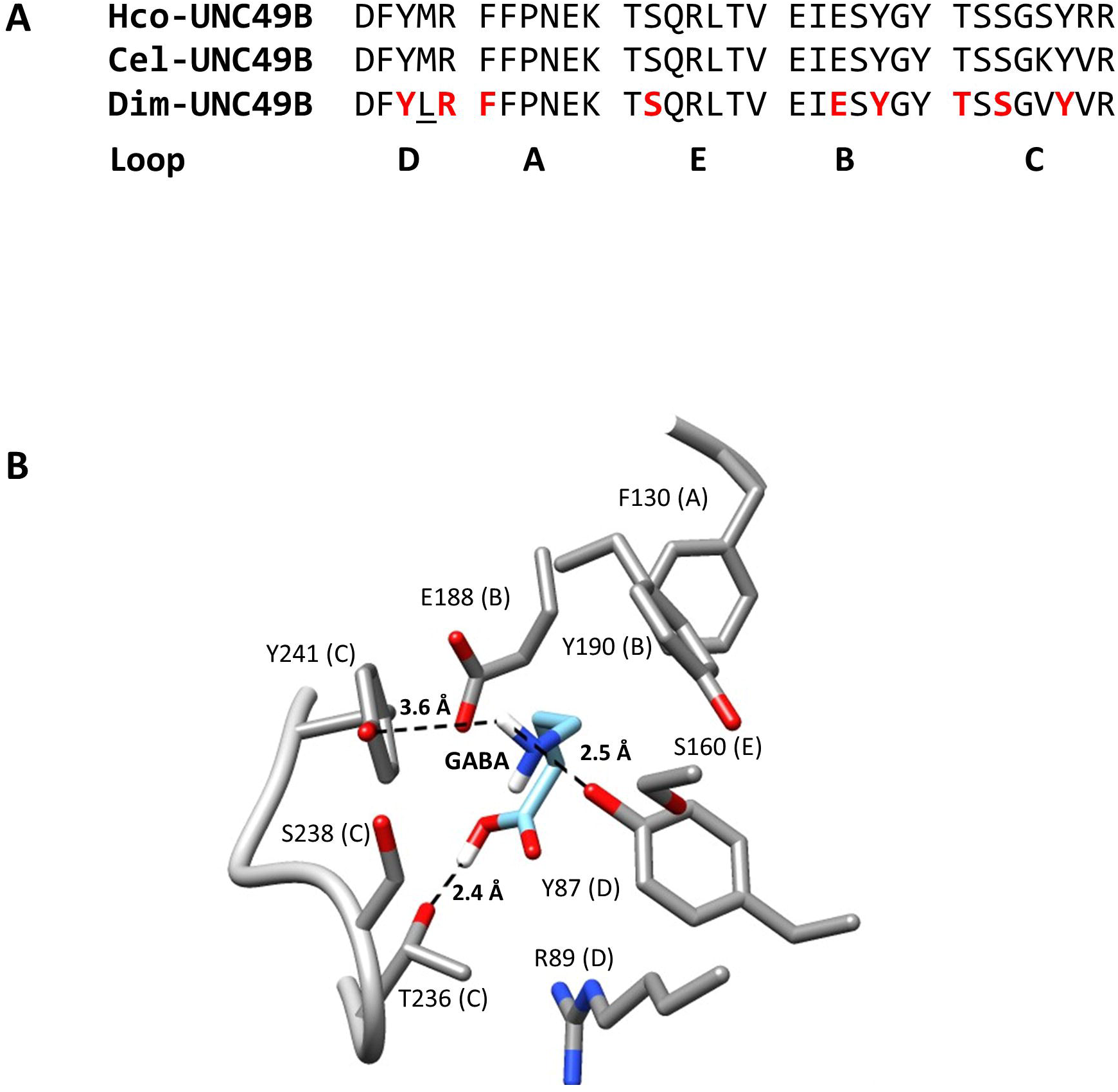
Dim-UNC-49 GABA binding site analysis. A. Alignments of the major binding loop sequences for UNC-49 from *D. immitis*, *H. contortus* and *C. elegans*. B. Major elements of the Dim-UNC-49 binding pocket with GABA docked.

### Functional characterization of the UNC-49 receptors

#### GABA sensitivity

The *dim-unc-49b* and *dim-unc-49c* sequences were reverse transcribed into cRNA through an *in vitro* transcription reaction for functional investigation in the *Xenopus* oocyte expression system. We first expressed Dim-UNC-49B and Dim-UNC-49C singly to undertake the reconstitution of homomeric channels. As seen in other UNC-49 receptor studies, Dim-UNC-49B gave rise to functional GABA-gated channels (Figure 3A). As recognized for Hco-UNC-49C and Cel-UNC-49C subunits (Bamber et al 1999; Siddiqui et al 2010), expression of Dim-UNC-49C alone did not form a channel that could respond to GABA (Supplemental C). Furthermore, the co-injection of oocytes with cRNA of both *unc-49b* and *unc-49c* also responded to GABA suggesting that a heteromeric channel was formed (Figure 3A).

**Figure 3:**
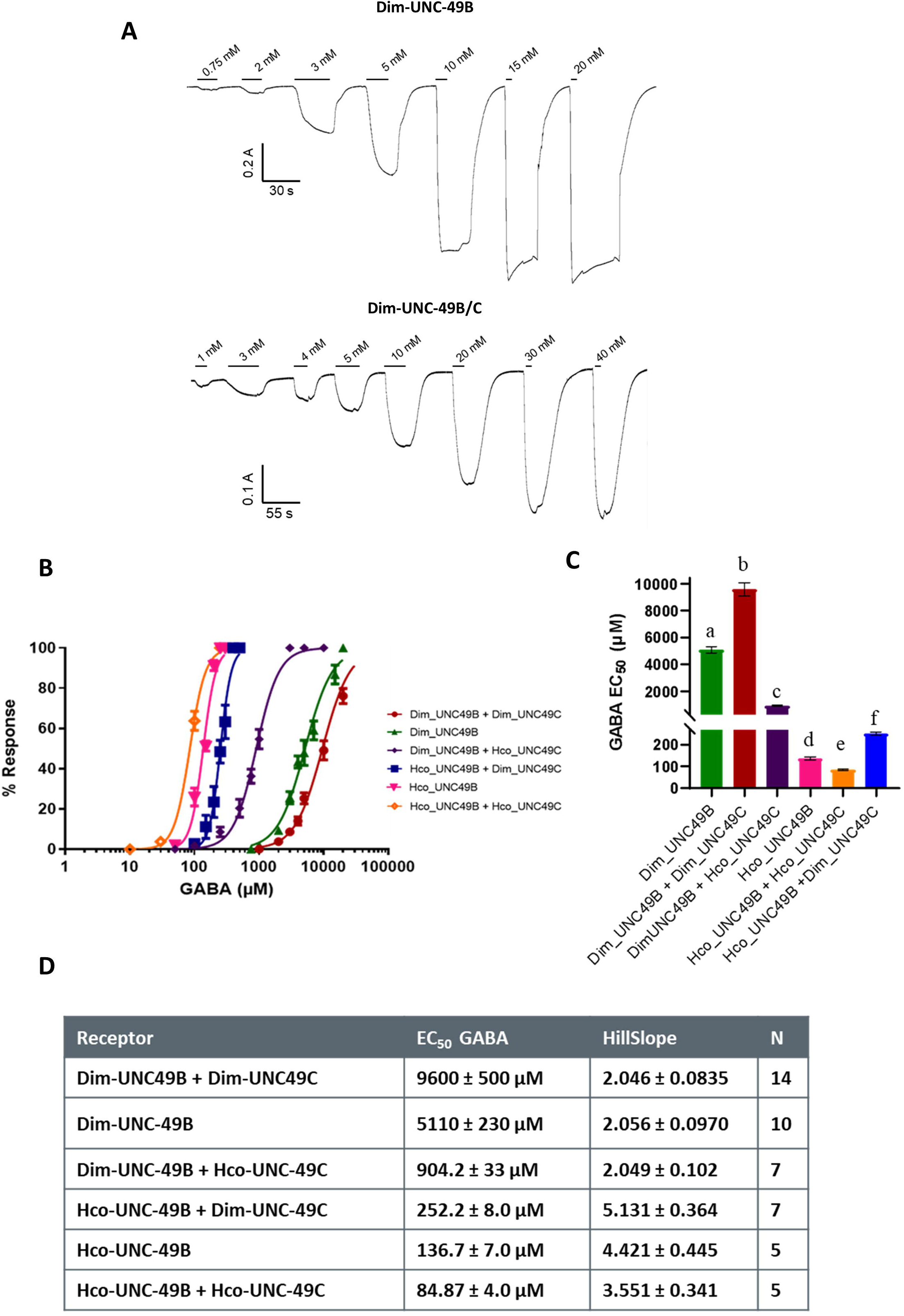
Dim-UNC-49 is a cys-loop GABA receptor. A. Representative traces of both the Dim-UNC-49 B homomeric channel and Dim-UNC-49 B/C heteromeric channel in response to increasing concentrations of GABA. (B) Dose response curves of all UNC-49 subunit combinations tested. (C) GABA EC_50_ values for all receptor combinations tested along with the statistical analysis. No significant difference (p≤0.05) between receptors identified by the same letter. Dim-UNC-49B receptors combinations have significantly higher EC_50_ when compared to Hco-UNC-49B receptors. In addition, Dim-UNC-49C significantly increases the EC_50_, whereas Hco-UNC-49C significantly decreases it in all receptor combinations tested. (D) EC_50_ and Hill Slope values obtained for all the receptors analyzed.

To further characterize the sensitivity of the *D. immitis* UNC-49 receptors to GABA and to confirm whether homomeric and heteromeric channels were formed in oocytes, we conducted dose-response experiments. This determined that GABA activates the homomeric Dim-UNC-49B channel with an EC_50_ of 5.11 ± 0.23 mM (n = 10) (Figure 3B and C). Heteromeric channels formed via co-expression with Dim-UNC-49C (Dim-UNC49BC) produced a channel with a lower sensitivity to GABA with an EC_50_ value of 9.60 ± 0.50 mM (n = 14). EC_50_ values between homomeric and heteromeric channels were statistically significant (Figure 3B and C). Since it appears that the Dim-UNC-49 homomeric and heteromeric channels have a much lower sensitivity than previous reports of the same channel in *C. elegans* (Bamber et al 1999) or *H. contortus* (Siddiqui et al 2010), we decided to reproduce the dose-response analysis on the *H. contortus* UNC-49 homomeric and heteromeric channels. When Hco-UNC-49B was expressed alone, the GABA concentration-response curve was characterized by an EC_50_ of 136.7 ± 7.0 µM (n = 5). When Hco-UNC-49B was co-expressed with Hco-UNC-49C, the EC_50_ decreased to 84.87 ± 4.0 µM (n = 5). The range and trend were similar to our previous reports (Siddiqui et al 2010). In addition, the use of the UNC-49B and C subunits from *H. contortus* allowed for a deeper investigation of the properties of the *D. immitis* receptors by conducting cross-species expression of the various subunits. Overall, we found that all subunits, regardless of whether they are *Dirofilaria* or *Haemonchus* were able to co-assemble with subunits from the other species which produced unique channels with statistically different EC_50_ values (Figure 3C). When Dim-UNC-49B was co-expressed with Hco-UNC-49C, it produced a channel with a GABA EC_50_ of 904.2 ± 33 µM which is reduced by 10-fold compared to the Dim-UNC-49B/C channel. Likewise, when Hco-UNC-49B was co-expressed with Dim-UNC-49C, the EC_50_ of GABA was 252.2 ± 8.0 µM. Comparisons of the EC_50_ and hill slopes of the various channels are found in Figure 3D. Firstly, all channels containing Dim-UNC-49B subunits (either homomeric or heteromeric channels) had EC_50_ values significantly higher from channels containing Hco-UNC-49B subunits. Secondly, heteromeric channels containing Dim-UNC-49C subunits had higher EC_50_ values compared to homomeric channels. The opposite was found in heteromeric channels containing Hco-UNC-49C subunits. Thirdly, homomeric or heteromeric channels containing Dim-UNC-49B subunits had significantly lower hill slopes compared to channels containing Hco-UNC-49B subunits suggesting that in our oocyte system UNC-49 receptors from *D. immitis* exhibit less cooperativity when activated by GABA (Figure 3D).

Current amplitudes were also analyzed between receptors that were measured on the same day. Noticeably, Dim-UNC-49B/C heteromeric channels exhibited lower current amplitudes compared to Dim-UNC-49B homomeric channels. We therefore examined this in more detail using several batches of oocytes. Interestingly, co-expression of Dim-UNC-49B with Dim-UNC-49C resulted in receptors which had significantly lower currents than that of the homomeric Dim-UNC-49B receptor (Figure 4A). Conversely, when Dim-UNC-49B was co-expressed with Hco-UNC-49C, the current amplitude was higher compared to the Dim-UNC-49B homomeric channel on two out of the three days analyzed (Figure 4B). This suggests that Dim-UNC-49C, through associating with Dim-UNC-49B is resulting in lower current amplitudes in the *Xenopus* expression system.

**Figure 4:**
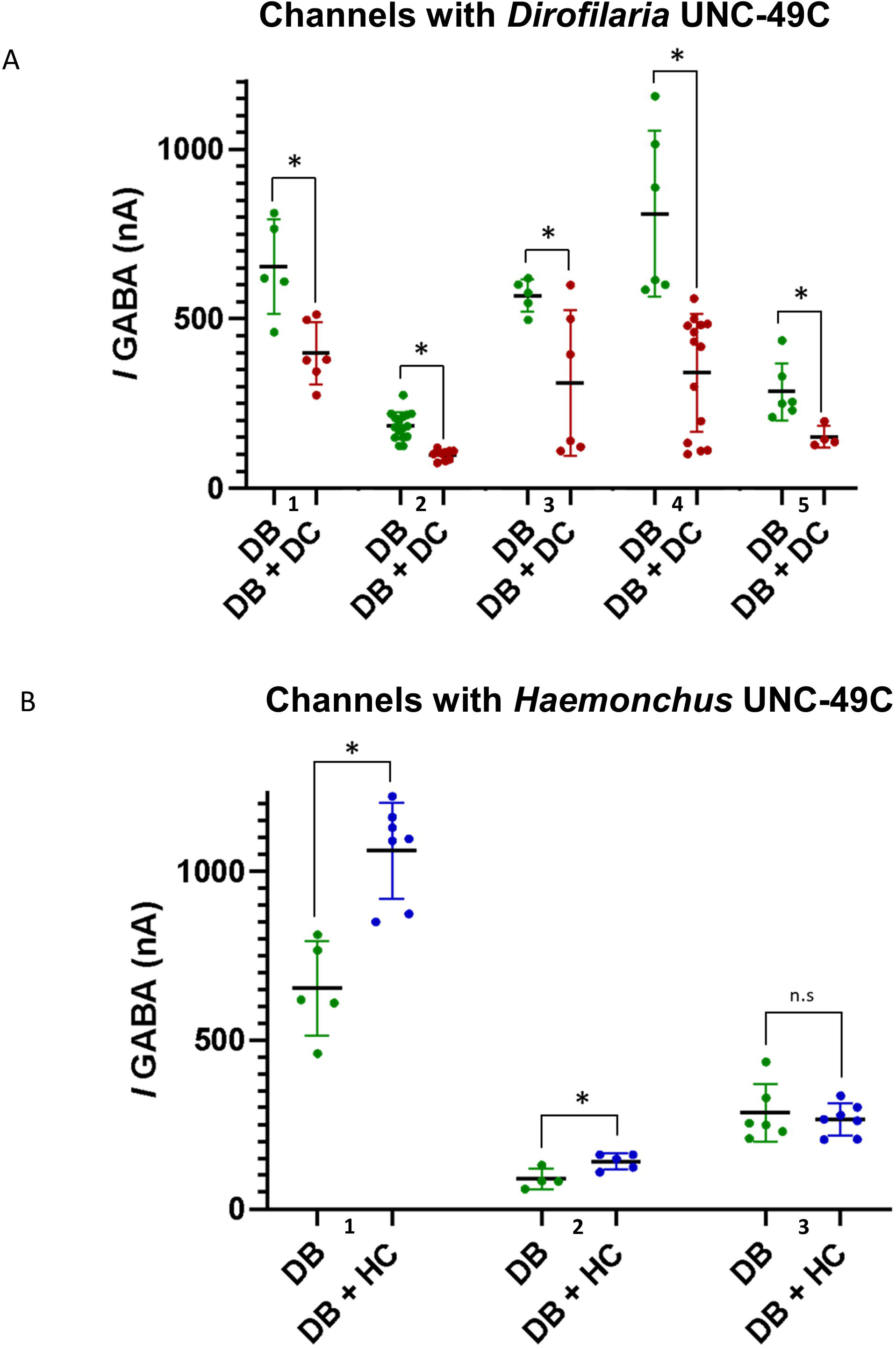
Comparison of the maximum amplitude measured between homomeric channels containing Dim*-*UNC-49B subunits and heteromeric channels containing either (A) Dim-UNC-49C or (B) Hco-UNC-49C. Concentrations of GABA used corresponded to the respective EC_50_ of the channel. Comparisons were performed on the same day and repeated using several batches of eggs (batches are indicated by the numbers 1 through 5 in Figure A and 1 through 3 in Figure B. Significance (p≤0.05) was determined through students t-test and displayed as asterisks. n.s indicates no significant difference (p > 0.05) between conditions. DB = Dim-UNC-49B homomeric channel; DB + DC = Dim-UNC-49BC heteromeric channel; DB + HC = Dim-UNC-49B/Hco-UNC-49C heteromeric channel. Each dot is an individual oocyte.

#### Picrotoxin (PTX) sensitivity

Previous studies have shown that the UNC-49 receptor was sensitive to the open channel blocker, PTX. Interestingly, it was shown the UNC-49B homomeric channels were more sensitive to PTX compared to heteromeric channels (Bamber et al 2003; Brown et al 2012). All subunit combinations were injected into *X. laevis* oocytes and the antagonistic response for PTX was measured by comparing the GABA response alone and in the presence of PTX. We first tested the PTX effect using a high concentration (250 µM). Here, it was found that unlike previous studies performed on *C. elegans* and *H. contortus* UNC-49 receptors, the Dim-UNC-49C subunit does not confer resistance to PTX. On the other hand, consistent with previous studies, Hco-UNC-49C subunit does confer PTX resistance (Figure 5A and B). To further investigate this phenomena, PTX dose response analysis was performed on all receptor combinations (Figure 6A). The Dim-UNC-49B homomeric channel was highly sensitive to PTX with an IC_50_ of 5.03 ± 0.23 μM and the Dim-UNC-49B/Dim-UNC-49C heteromeric channel was also highly sensitive to PTX with a similar IC_50_ of 4.15 ± 0.50 μM (Figure 6B and C). This showed that Dim-UNC-49C, while able to associate with UNC-49B did not result in a PTX-resistant receptor. This was confirmed when Dim-UNC-49C was co-expressed with Hco-UNC-49B. Here there was no significant difference between PTX IC_50_ for the Hco-UNC-49B/Dim-UNC-49C heteromeric channel (IC_50_ of 16.75 ± 3.15 μM) and the Hco-UNC-49B homomeric channel (IC_50_ of 21.4 ± 5.82 μM). A different effect was observed when we combined the Dim-UNC-49B subunit with Hco-UNC-49C. This heteromeric was much more resistant to PTX (IC_50_ of 268 ± 23 μM) which was consistent with our previous observation that Hco-UNC-49C directly causes PTX resistance in heteromeric channels (Brown et al 2012). The M2 domain has been shown to play a direct role in resistance to PTX. Alignment of the M2 domains of the UNC-49B subunits of *H. contortus, C. elegans* and *D. immitis* show high conservation between species (Figure 6D). On the other hand, there is more variability in the amino acids of the M2 domain of UNC-49C, with *D. immitis* showing the most variability compared to both *H. contortus* and *C. elegans*.

**Figure 5:**
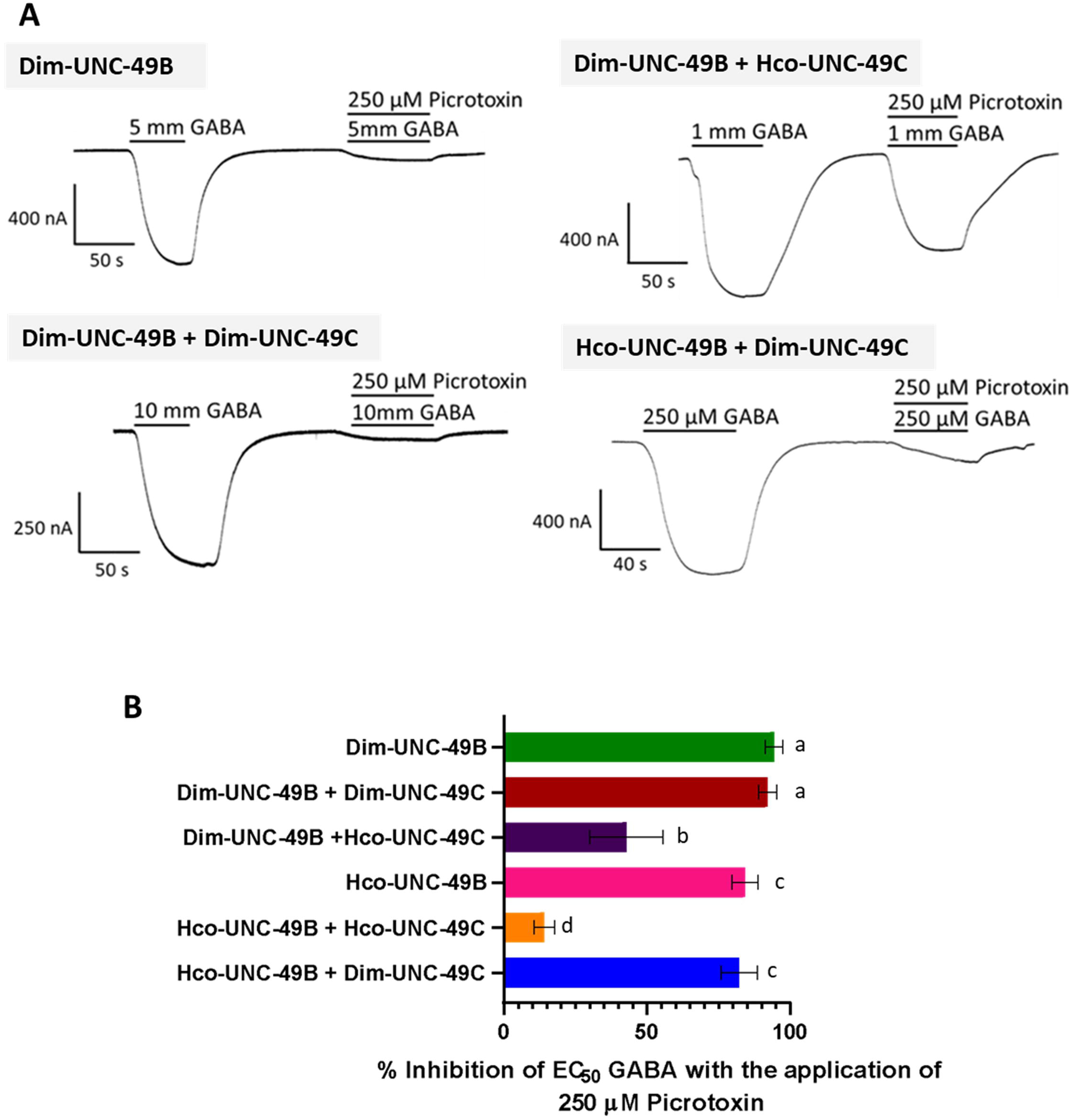
(A) Representative electrophysiological traces showing the effect maximal PTX concentration (250 μM) on EC_50_ GABA. Single line represents EC_50_ GABA application alone, double line represents EC_50_ GABA with 250 μM of PTX. (B) Maximum EC_50_ GABA inhibition, with statistical analysis. Groups with the same letter are not significantly different (*p* > 0.05) and groups with different letters are significantly different (*p* < 0.05).

**Figure 6:**
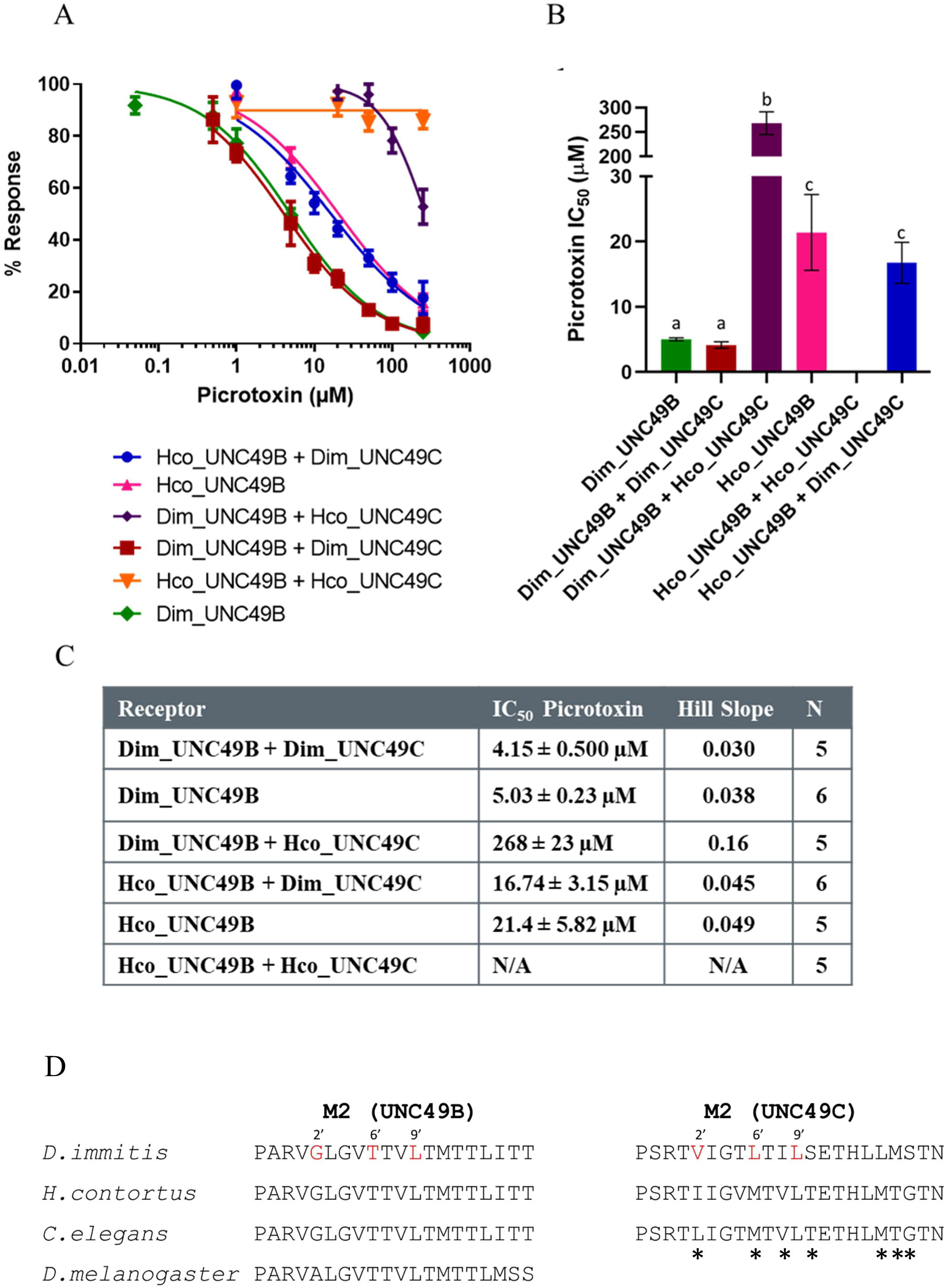
(A) PTX dose-dependent inhibition of GABA EC_50_ response for all receptor combinations tested. (B) Comparisons of IC_50_ values for PTX for all receptors analyzed. No significant difference (p ≤ 0.05) between receptors identified by the same letter. (C) Summary of the IC_50_ data, the hill slope and number of replicates (N). (D) Alignment of the M2 region of Dim-UNC-49 with other PTX sensitive receptors. Numbers 2’, 6’ and 9’ have been identified to be important for PTX sensitivity. * indicates residues that are variable when comparing the UNC-49C subunit between *D. immitis*, *H. contortus* and *C. elegans*.

## Discussion

*D. immitis* is an important parasite in many parts of the world and is the focus of ongoing research into the discovery of novel heartworm preventatives (Mwacalimba et al 2024). Despite this, information on the properties of cys-loop receptors in this organism is extremely lacking. The last paper published on the discovery of novel cys-loop receptors was published in 2004 and reported the cloning of a novel glutamate-gated chloride channel (GluCl) which is the target of the anthelmintic ivermectin (Yates and Wolstenholme 2004). Therefore, the paper presented here is only the second report of the discovery and characterization of a novel cys-loop receptor in this important pathogen.

There is now a growing body of evidence that GABA receptors play an important role in parasitic nematode motility. The original studies on GABA receptors in parasitic nematodes were performed on the whole worm body wall muscle from the parasites *Ascaris lumbricoides* and *A. suum* (Del Castillo et al. 1963, 1964a,b; Holden-Dye et al. 1988, 1989; Holden-Dye and Walker 1990; Martin 1980, 1985; Martin et al. 1991). Electrophysiological analysis of these muscle receptors determined the presence of a pharmacologically unique GABA receptor involved in muscle contraction and movement. These results formed the initial evidence that GABA receptors found in parasitic nematodes could make promising targets for novel antiparasitics.

Subsequently, research using the model nematode *C. elegans* led to the discovery of a sequence of a cys-loop GABA receptor encoded for by the *unc-49* gene (Bamber et al. 1999). This receptor was found at neuromuscular junctions and was responsible for the regular sinusoidal movement of worms. Subsequent research on the related parasitic nematode *H. contortus* led to the discovery of orthologues of the *unc-49* gene (Siddiqui et al 2010). These receptors were found to have a unique pharmacology that shared similarity to the profile of the original muscle GABA receptors in *Ascaris* (Kaji et al 2015). Namely, the relative sensitivity of the UNC-49 receptor to compounds such as muscimol and GABOB and its insensitivity to sulphonated agonists were similar to the *Ascaris* receptor (Kaji et al 2015). It’s important to note that UNC-49 GABA receptors are not direct orthologues of mammalian GABA receptors but rather share more similarity to insect RDL GABA receptors (Bamber et al. 2003). As the genomes of more parasitic nematodes become sequenced, we are noticing that genes encoding UNC-49 GABA channels are found throughout the nematode phylum (Accardi et al 2012). As for *D. immitis* we have now confirmed that the *unc-49* gene is also present and like both *C. elegans* and *H. contortus* there is differential splicing producing individual subunits (UNC-49B and C).

While *unc-49* receptor genes are present throughout nematoda, the functional properties of the channels expressed in oocytes differ depending on the species. For example, in the *C. elegans* UNC-49 receptor, the UNC-49C subunit appears to modulate the function of the heteromeric channel expressed in oocytes where it increases the EC_50_ compared to the homomeric channel. In addition, when expressed in HEK cells, single-channel conductance of UNC-49B/C heteromeric channels was reduced by 20% compared to the UNC-49B homomeric channel (Bamber et al 1999). The Dim-UNC-49C subunit also appears to have a similar effect on the Dim-UNC-49BC heteromeric channel and is less sensitive to GABA compared to the homomeric channel. This was confirmed when we co-expressed the *H. contortus* UNC-49B with *D. immitis* UNC-49C where the heteromeric channel was less sensitive to GABA compared to the *H. contortus* UNC-49B homomeric channel. Interestingly, the opposite was found when we analyzed *H. contortus* UNC-49C containing channels. We find that the heteromeric channels containing *H. contortus* UNC-49C are more sensitive compared to the homomeric channel regardless of whether it was co-expressed with an UNC-49B subunit from *H. contortus* or *Dirofilaria* (Siddiqui et al 2010; present study). Not only did the *H. contortus* UNC-49C subunit increase sensitivity of heteromeric channels, but it also increased the current observed in the oocyte system. So taken together, our data suggest that Dim-UNC-49C is having an effect not only on the GABA sensitivity of heteromeric channels but on the amount of current observed.

Interestingly, studies on epilepsy disorders have identified an α1 GABA receptor variant that has a G251S amino acid change immediately before membrane spanning domain 1 that is associated with Dravet Syndrome (Carvill et al 2014). When this variant was expressed in *Xenopus* oocytes, there was a significant reduction in both GABA sensitivity and current (Carvill et al 2014). Strikingly, a similar mutation, G254D, introduced in the analogous position in the UNC-49B subunit in *C. elegans* significantly reduces movement (Gadhia et al 2024). In the *D. immitis* UNC-49C subunit there is a naturally occurring serine at position 256 where both *H. contortus* and *C. elegans* UNC-49C have a glycine (Figure 8). It would be interesting to examine this residue position in Dim-UNC-49C using both electrophysiology and expression in *C. elegans* mutants to determine whether it plays any functional role.

**Figure 7:**
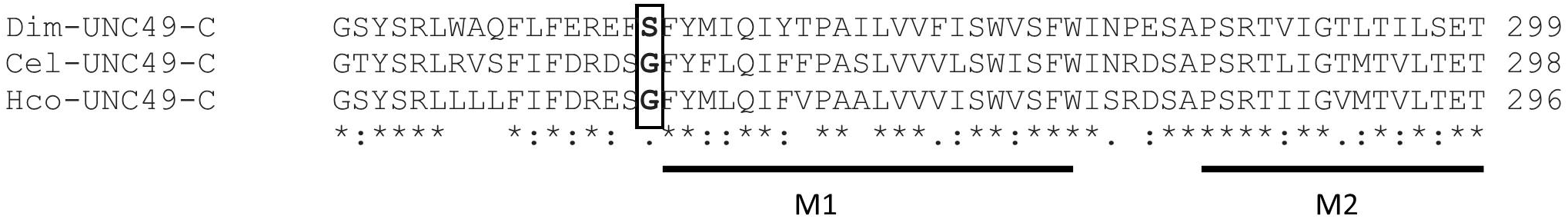
Alignment of sequence near the M1 and M2 domains of UNC-49C subunits. Box indicates the position that has been shown to be associated with Dravet Syndrome in the human alpha1 subunit and movement in the *C. elegans* UNC-49B subunit.

It has been previously reported that both the *H. contortus* and *C. elegans* UNC-49C subunits confers resistance to PTX block (Bamber et al 2003; Brown et al 2012). This was also confirmed when we co-expressed *D. immitis* UNC-49B with *H. contortus* UNC-49C. However, the *D. immitis* UNC-49C subunit does not confer resistance when associated with either the *D. immitis* or *H. contortus* UNC-49B subunit. Previous research on the RDL receptor from insects have revealed several residues in the M2 region important for PTX binding. These include alanine (2’), threonine (6’) and leucine (9’) (Figure 6D). Modeling studies on the RDL receptor have shown that the 6’T forms a hydrogen bond with PTX with 2’A and 9’L contributing hydrophobic interactions. The analogous residues in the M2 domain of UNC-49B in nematodes is indicated in figure 6D. Here, it shows that although nematodes have a G instead of A at 2’ the other residues important for PTX-binding are present. Consistent with this, we observed that the UNC-49B homomeric channel is PTX sensitive. It has been suggested that UNC-49C provides PTX resistance through the loss of a threonine at 6’ which is replaced by a methionine in both *H. contortus* and *C. elegans* UNC-49C (Bamber et al 2003). In *D. immitis* UNC-49C, the threonine at 6’ is replaced by a leucine (not methionine) and therefore we could only speculate about the role of this amino-acid in PTX sensitivity. In addition, there are several other amino acid differences in the M2 domain of Dim-UNC-49C compared to both the *C. elegans* and *H. contortus* UNC-49C subunit. Clearly, further experiments are required to decipher the PTX sensitivity/resistance determinants in UNC-49 subunits from nematodes.

Overall, we have found that like other nematodes, *D. immitis* also exhibit UNC-49 GABA receptors. However, it appears that compared to similar receptors found in the parasitic nematode *H. contortus*, there are clear functional differences. However, given that the pharmacology of the UNC-49 receptor is different compared to mammalian GABA receptors (Kaji et al 2015) it may be possible to design a drug that targets nematode GABA receptors with minimal effects on similar receptors in the mammalian host. Future research is required to establish whether UNC49 GABA receptors are potential targets for novel nematocides.

