## Supplemental Figures for "The *Dirofilaria immitis unc-49* gene encodes a pharmacologically unique cys-loop GABA receptor"

```
Dim_UNC49_B      MHLISFFFIIYCTVYNYTLGYIEYPSSKTFNTTAISDILNRLVDKSTYDKRLRPKYGAEP 60
Cel_UNC49_B      MARPFTLIVLLSAHLCLH-VVVTQDEDSHINTQLLSSVLDRLTNRTTYDKRLRPRYGEKP 59
Hco_UNC49_B      -MRISDLFLFATTATFYH-VSRAQEDDIIVDTSQLSTVLDRLTNKTIYDKRLRPRYGDKP 58
                       ::::  :            ..   .:*   :*  :*:**.:::  *******:**  :*
                   Signal Peptide

Dim_UNC49_B      VDVGITIHVSSISAVSEVDMDFTIDFYLRQTWQDPRLAFGDMFYGYQQGKIESLTVGVDY 120
Cel_UNC49_B      VDVGITIHVSSISAVSEVDMDFTLDFYMRQTWQDPRLAFGSLDLGLS-KEIDSLTVGVDY 118
Hco_UNC49_B      VDVGITIHVSSISAVSEVDMDFTLDFYMRQTWQDPRLAFGTLDLGIA-KQITSLTVGVDY 117
                 ***********************:***:************  :  *     :* ********
                                          Loop D

Dim_UNC49_B      LDKLWKPDTFFPNEKKSFFHTATTHNSFLRIDPDGTVFTSQRLTVTATCPMKLQLFPMDS 180
Cel_UNC49_B      LDRLWKPDTFFPNEKKSFFHLATTHNSFLRIEGDGTVYTSQRLTVTATCPMDLKLFPMDS 178
Hco_UNC49_B      LDRLWKPDTFFPNEKKSFFHLATTHNSFLRIDSDGTVYTSQRLTVTATCPMKLQLFPMDS 177
                 **:***************** **********:  ****:*************.*:******
                           Loop A          Loop E

Dim_UNC49_B      QKCKLEIESYGYTTADIDLFWGKDRKDQGQ-VVGFENISLPQFKPVGYRVNVTRAITSSG 239
Cel_UNC49_B      QHCKLEIESYGYETKDIDYYWGKKRTDLEITAVKFDTFQLPQFQPTLYFVNTTKAETSSG 238
Hco_UNC49_B      QRCKLEIESYGYTMADIDYFWGRQRSDPRQSAVVFGNFMLPQFKQTCYHVNYTQATTSSG 237
                 *:********* *   *** :**:.*.*     .* *  .: ****:  .  *  **  *:* ****
                      Loop B                                      Loop C

Dim_UNC49_B      VYVRLYFEVLLARNLGFYLMDIIIPSMLIVTISWVSFWLNREASPARVGLGVTTVLTMTT 299
Cel_UNC49_B      KYVRLALEVILVRNMGFYTMNIVIPSILIVTISWVSFWLNREASPARVGLGVTTVLTMTT 298
Hco_UNC49_B      SYRRLYFEILLVRNMGFYSMNIVIPSMLIVTISWVSFWLNREASPARVGLGVTTVLTMTT 297
                  * **  :*::*.**:***  *:*:***:*********************************
                                      M1                        M2

Dim_UNC49_B      LITTTNNAMPKVSYIKGLDVFLNFCFVMVFASLIEYAVVSYMNKKLVQRREKRRKQVEQG 359
Cel_UNC49_B      LITTTNNSMPKVSYVKGLDVFLNFCFVMVFASLLEYAIVSYMNKRLVLRREKRRKAAEQQ 358
Hco_UNC49_B      LITTTNNSMPKVSYIKGLDVFLNFCFVMVFASLVEYAVVSYMNKRIALRREKRRRQAEQQ 357
                 *******:******:*****************:***:******::.  ******:  .**
                                    M3

Dim_UNC49_B      IPVEMPMYNQLTTNAETKMQMDTFLG-SSGQNP-----NLSVSEALIPECDCRTIPLIQN 413
Cel_UNC49_B      QRNEMPMFNASPKAAN----------------------NNNPLMEIPENCDCRTIPMMQH 396
Hco_UNC49_B      QRNEVPMFSNPVTPKQPNNNVYEMAMISQNSTPAKTFVPHSQLMEIPVDCDCRTIPLIQH 417
                    *:**:.    .   :                           .     :   :*******::*:

Dim_UNC49_B      PRLVAD--HTLWPAPFMRPKRASRTC-RSVTPSKIDKFSRFLFPMLFLAFNLLYWTIMTV 470
Cel_UNC49_B      PRLVTDGAHTLWPAPFARPKKASKTCCQRWTPAKIDKLSRYGFPLSFSIFNIVYWLYMKY 456
Hco_UNC49_B      PRLVADGTHTMWPAPFGKPKKASKTC-RNVTPAKIDKCSRYLFPLLFSAFNVVYWTIMTV 476
                 ****:*  **:***** :**:**:** :  **:**** **: **: *  **::**  *.
                                                    M4

Dim_UNC49_B      LSSFVDTSDYTLFD-------- 484
Cel_UNC49_B      LSLNSSDKIQENDKWQQIH--- 475
Hco_UNC49_B      LSSIAED--LSKDEWVPIVIED
```

Supplemental A: Protein sequence alignment of *D. immitis* UNC-49B with *C. elegans* UNC-49B*,* and *H. contortus* UNC-49B. Asterisks (*) indicate identical amino acids and (:) represents similar amino acids. The binding loops, and trans-membrane domains (TM) are displayed. Stars indicate residues where there are variations from the *H. contortus* receptor, which have been shown previously to be important for GABA sensitivity (Accardi and Forrester 2011).

```
Dim_UNC49_C      MHLISFFFIIYCTVYNYTLGYIEYPSSKTFNTTAISDILNRLVDKSTYDKRLRPKYGAEP 60
Cel_UNC49_C      MARPFTLIVLLSAHLCLH-VVVTQDEDSHINTQLLSSVLDRLTNRTTYDKRLRPRYGEKP 59
Hco_UNC49_C      -MRISDLFLFATTATFYH-VSRAQEDDIIVDTSQLSTVLDRLTNKTIYDKRLRPRYGDKP 58
                       ::::  :            ..  .:*  :* :*:**.::: *******:** :*
                   Signal Peptide

Dim_UNC49_C      VDVGITIHVSSISAVSEVDMDFTIDFYLRQTWQDPRLAFGDMFYGYQQGKIESLTVGVDY 120
Cel_UNC49_C      VDVGITIHVSSISAVSEVDMDFTLDFYMRQTWQDPRLAFGSLDLGLS-KEIDSLTVGVDY 118
Hco_UNC49_C      VDVGITIHVSSISAVSEVDMDFTLDFYMRQTWQDPRLAFGTLDLGIA-KQITSLTVGVDY 117
                 ***********************:***:************ :  *    :* ********
                                           Loop D

Dim_UNC49_C      LDKLWKPDTFFPNEKKSFFHTATTHNSFLRIDPDGTVFTSQRLTVTATCPMKLQLFPMDS 180
Cel_UNC49_C      LDRLWKPDTFFPNEKKSFFHLATTHNSFLRIEGDGTVYTSQRLTVTATCPMDLKLFPMDS 178
Hco_UNC49_C      LDRLWKPDTFFPNEKKSFFHLATTHNSFLRIDSDGTVYTSQRLTVTATCPMKLQLFPMDS 177
                 **:***************** **********: ****:*************.*:******
                         Loop A              Loop E

Dim_UNC49_C      QKCKLEIESYGYTVATIHYHWCQLKDIDCKTAVKVE-PFELTSYRFANLCINKTIATTSS 239
Cel_UNC49_C      QHCKLEIESYAYSTAEIEYKWCTSKEPNCSTAVKADANIELSSYKFTKICQKRTLASTSS 238
Hco_UNC49_C      QRCKLEIESYGYSTAAIEYHWCGAANPNCETAVVAD-DVELPSYRFAKVCIDRTMATTAS 236
                 *:********.*:.* *.*:**   : :*.*** .:  .** **:*:::* .:*:*:*:*
                     Loop B                                         Loop C

Dim_UNC49_C      GSYSRLWAQFLFEREFSFYMIQIYTPAILVVFISWVSFWINPESAPSRTVIGTLTILSET 299
Cel_UNC49_C      GTYSRLRVSFIFDRDSGFYFLQIFFPASLVVVLSWISFWINRDSAPSRTLIGTMTVLTET 298
Hco_UNC49_C      GSYSRLLLLFIFDRESGFYMLQIFVPAALVVVISWVSFWISRDSAPSRTIIGVMTVLTET 296
                 *:****   *:*:*: .**::**: ** ***.:**:****. :******:**.:*:*:**
                                    M1                     M2

Dim_UNC49_C      HLLMSTNKRLPPVSYIKAVDVYLGFCYLNVALALIEFATVAYTKKKYEDRKRNKSKNVFE 359
Cel_UNC49_C      HLMTGTNRRLPPVAYVKAVDVFLGFCYLLVILALIEYACVAYSKKKNEDRRRREKKT--E 356
Hco_UNC49_C      HLMTGTNRRLPPVAYVKAVDVYLGFCYLLVVLALIEYACVAYSKKKNDDRRRREKKS--E 354
                 **: .**:*****:*:*****:****** * *****:* ***:*** :**:*.:.*.  *
                                      M3

Dim_UNC49_C      LAPQVQAPDLLRDVRIDVCTCDQNVSVFLVDKKSPKCNNCCIHSRLDLIARIAFPISFLM 419
Cel_UNC49_C      HKPAPPTPDILHDVRLAECTCNAAPTSIIAVIKQ-SNRFCVSHSHIDIVSRAAFPLVFIL 415
Hco_UNC49_C      HKPTPPTPDLLNDARLAECTCNAAPTSIIAVIKQ-PNRFCIRHSHIDIASRVVFPCTFLF 413
                   *   :**:*.*.*:  ***:   : ::.  *.   . *  **::*: :* .**  *::
                                                                M4

Dim_UNC49_C      FNFIYWTVLLSLSGWKIYGDNTEQRAC------ 446
Cel_UNC49_C      FNTLFWLILLYKSKRLPYISEHEGDRCDAPDLH 448
Hco_UNC49_C      FNMLFWVVLLAKAKRLPYFSTAAVPRC------ 440
```

Supplemental B: Protein sequence alignment of *D. immitis* UNC-49C with *C. elegans* UNC-49C*,* and *H. contortus* UNC-49C. Asterisks (*) indicate identical amino acids and (:) represents similar amino acids. The binding loops, and trans-membrane domains (TM) are displayed.

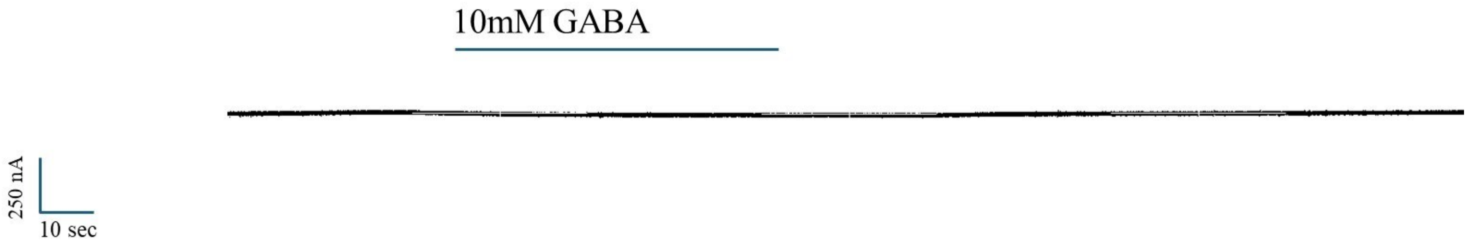


Supplemental C. UNC-49C expression in oocytes showing no response to GABA.
